# Omics-based Hybrid Prediction in Maize

**DOI:** 10.1101/134668

**Authors:** Matthias Westhues, Tobias A. Schrag, Claas Heuer, Georg Thaller, H. Friedrich Utz, Wolfgang Schipprack, Alexander Thiemann, Felix Seifert, Anita Ehret, Armin Schlereth, Mark Stitt, Zoran Nikoloski, Lothar Willmitzer, Chris C. Schön, Stefan Scholten, Albrecht E. Melchinger

## Abstract

Accurate prediction of traits with complex genetic architecture is crucial for selecting superior candidates in animal and plant breeding and for guiding decisions in personalized medicine. Whole-genome prediction (WGP) has revolutionized these areas but has inherent limitations in incorporating intricate epistatic interactions. Downstream “omics” data are expected to integrate interactions within and between different biological strata and provide the opportunity to improve trait prediction. Yet, predicting traits from parents to progeny has not been addressed by a combination of “omics” data. Here, we evaluate several “omics” predictors — genomic, transcriptomic and metabolic data — measured on parent lines at early developmental stages, and demonstrate that the integration of transcriptomic with genomic data leads to higher success rates in the correct prediction of untested hybrid combinations in maize. Despite the high predictive ability of genomic data, transcriptomic data alone outperformed them and other predictors for the most complex heterotic trait, dry matter yield. An eQTL analysis revealed that transcriptomic data integrate genomic information from both, adjacent and distant sites relative to the expressed genes. Together, these findings suggest that downstream predictors capture physiological epistasis that is transmitted from parents to their hybrid offspring. We conclude that the use of downstream “omics” data in prediction can exploit important information beyond structural genomics for leveraging the efficiency of hybrid breeding.

**Key message:** Complementing genomic data with other “omics” predictors can increase the probability of success for predicting the best hybrid combinations using complex agronomic traits.

**Conflict of Interest:** The authors declare that they have no conflict of interest.

## Introduction

Hybrid breeding, which entails crossing of lines from two genetically distant germplasm collections — called heterotic groups (Melchinger and Gumber, 1998) — has emerged as a prime strategy to meet demands for a sustainable intensification of agricultural production (Duvick, 2005). However, unlocking the full potential of hybrid breeding requires accurate prediction methods to efficiently identify the superior candidates out of the millions of possible hybrids that could potentially be produced in each cycle of an ordinary-sized breeding program. With the advent of the doubled haploid (DH) technology (Wedzony et al., 2009) this prediction problem has become even more challenging because, based on breeder’s experience, the vast majority (*≈* 90%) of competing lines in each heterotic group are “new” lines without any phenotypic records on hybrid progeny from previous breeding cycles. Consequently, among all hybrid combinations possible between lines from two heterotic groups, about 81% are T0 hybrids, 18% are T1 hybrids and 1% are T2 hybrids having zero, one or two parents, respectively, that have been previously tested in other hybrid combinations. Preselection of a few hundred of the most favorable hybrids with high success rate could significantly reduce the labor-intensive and time-consuming field-testing (Kadam et al., 2016, Xu et al., 2016). This could greatly impact the efficiency of hybrid breeding and boost the annual selection gain (Longin, Mi and Würschum, 2015).

Whereas yield and other heterotic traits of hybrids are generally poorly predicted by the performance of their parent lines (Melchinger and Gumber, 1998), WGP has emerged as a major tool for tackling this challenge (Massman et al., 2013, Technow et al., 2014). Nevertheless, there is evidence that, even with complete sequence information, genomic prediction may not capture complex interactions between genes and downstream regulation, which act through the entire cascade from genotype to phenotype (Dalchau et al., 2011, Zhu et al., 2012, Rudd et al., 2015, Ritchie, Holzinger, Li, Pendergrass and Kim, 2015). Most studies have evaluated predictive ability by looking at only one kind of endophenotype (intermediaries between genotype and phenotype (Gottesman and Gould, 2003, Mackay, Stone and Ayroles, 2009) such as the transcriptome (Swanson-Wagner et al., 2006, Zenke-Philippi et al., 2016, Xu et al., 2016) or the metabolome (Riedelsheimer et al., 2012, Xu et al., 2016, Dan et al., 2016). The integration of different endophenotypic and genomic data is expected to reflect more closely the variability across genotypes than genomic data alone (Mackay, Stone and Ayroles, 2009, Patti, Yanes and Siuzdak, 2012, Civelek and Lusis, 2014). Two recent studies that integrated multiple biological strata in predicting breast cancer risk (Vazquez et al., 2016) and performance of maize inbred lines (Guo et al., 2016), respectively, demonstrated the benefit of this strategy. However, unlike forecasting clinical or agronomic traits from endophenotypes of the same genotype, hybrid breeding requires the prediction of the genotypic values (GV) of hybrid progeny based on parental information. To achieve this objective, we used the BLUP approach — originally developed in animal breeding (Henderson, 1984) — for the more complex setting of hybrids between parents from two heterotic groups (Bernardo, 1996, Massman et al., 2013). Here, we measured endophenotypes of parent lines to forecast the GV of T0, T1 and T2 hybrid progeny by using prediction equations trained with “omics” information from other parent lines and phenotypic information on their hybrid offspring.

## Materials and Methods

### Genetic material and phenotyping

The entire genetic material consisted of a set of 1,536 hybrids, denoted as *H*_*Tot*_, produced in 16 factorial mating designs between 142 Dent and 103 Flint lines from the maize breeding program at the University of Hohenheim, on which agronomic data for silage maize production, as well as pedigree and genomic data were available. A subset of this material, albeit from different trials, has been used for genomic prediction of traits related to grain maize production (Technow et al., 2014). For hybrid prediction, we used a core set *H ⊂ H*_Tot_ of 617 hybrids, produced in six factorials with hybrid sets 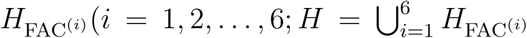 from crosses between 57 Dent and 41 Flint inbred lines, denoted as *D* = {1, 2, *…,* 57} and *F* = {1, 2, *…,* 41} (File S1). All hybrids were evaluated in field experiments at three or more agro-ecologically diverse locations across Germany. In the trials of each factorial, which included at least five common check genotypes, the entries were randomized in *α* lattice designs and planted in 2-row plots. Dry matter yield (DMY, *t/ha*) and dry matter content (DMC, %) of whole-plant aboveground biomass were determined by established procedures (Riedelsheimer et al., 2012). For quality traits, contents of fiber (ADF, %), fat (FAT, %_0_), protein (PRO, %_0_), starch (STA, %), and sugars (SUG, %_0_) in dry matter were measured in the harvested plant material using calibrated near-infrared spectroscopy (NIRS; Grieder et al. (2011), File S1).

### Pedigree-based relationship coefficients

Coancestry coefficients were calculated using SAS (version 9.4, SAS Institute) for all possible pairs of lines in each heterotic group according to established rules (Falconer and Mackay, 1996) under the following assumptions (Cox, Murphy and Rodgers, 1986): (i) all lines in a pedigree are genetically homogeneous and homozygous, (ii) pairs of genotypes with no known common parentage are unrelated, and (iii) a line derived from a cross or backcross obtained a proportional fraction of the genome from each parent, as expected under Mendelian inheritance in the absence of selection.

### Genotyping

Genotyping of all inbred lines was performed with the Illumina SNP chip MaizeSNP50 (Ganal et al., 2011). After performing a commonly used quality check (Technow et al., 2014) and imputation of missing data (Browning and Browning, 2009), a total of 21,565 polymorphic SNPs was available and used for all further analyses.

### Metabolite profiling

Seedlings of all parental inbred lines were grown under controlled conditions inside climate chambers to quantify the metabolite profiles of their roots 3.5 days after sowing, as detailed by de Abreu e Lima et al. (2017). The experiment was laid out as a randomized incomplete block design with replicated germination boxes. For leaf metabolic profiles (known metabolites and unannotated chromatographic peaks), a field experiment was carried out in an *α* lattice design with two replications at one location in southern Germany in the spring of 2012. Excision of leaves at the third leaf stage was performed according to an established protocol (Riedelsheimer et al., 2012) 28 days after sowing, in the afternoon of a cloudy day, and finalized within 45 minutes for the entire experiment. For both profiling procedures, all material was transferred directly into containers with dry ice and then into liquid nitrogen to quench metabolic activity.

### Transcriptome profiling

For transcriptome profiling, five seeds per parent line were taken from the same seed lot as used for metabolite profiling and laid out inside a climate chamber in a randomized complete block design with five replications. Seedlings were sampled seven days after sowing, snap-frozen in liquid nitrogen, and stored at -80°C until use. Prior to mRNA extraction, roots from all replicates of a genotype were pooled and homogenized. A custom 2K-microarray (GPL22267) was assembled from a subset of the 47K maize oligonucleotide array (GPL6438). Two-color hybridizations were carried out separately for each of the six factorials using interwoven loop designs (Kerr and Churchill, 2001). The average number of shared genotypes between factorials was 4.5 and ranged from 2 to 10.

### Statistical analysis of agronomic traits

Agronomic data were analyzed in two stages, following Technow et al. (2014) by accounting for year, location, field replication, block and genotype effects as well as their interactions (File S1). In the first stage, and separately for each environment, best linear unbiased estimates (BLUEs) of the *α* designs were computed for every hybrid using REML-based linear mixed-model analyses. In the second stage, BLUEs were computed for all hybrids in *H*_*Tot*_. The BLUEs of hybrids in the core set *H* served as response variables in our hybrid prediction models and cross-validation routines. For all predictions we used computationally efficient best linear unbiased predictor (BLUP) models, which have the same properties as those of a selection index because we previously accounted for fixed effects (Mrode (2014), pp. 34, 311, 312). For general and specific combining abilities (GCA and SCA) of parent lines, we used ASReml (Butler et al., 2009) to compute best linear unbiased predictors (BLUPs), variance components 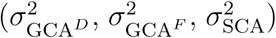 and entry-mean heritabilities (*H*^2^) of all hybrids in *H*_*Tot*_, treating all effects in the model as random. The covariance matrices of the GCA and SCA effects were defined by multiplying the variance components with their respective genomic relationship matrices (File S1).

### Statistical analysis of endophenotypes

Raw data were normalized using established procedures for metabolites (van den Berg et al., 2006) and transcripts (Smyth and Speed, 2003, Ritchie et al., 2007). From these data, we obtained BLUEs for metabolite levels and transcript abundance of each line using REML-based mixed-model analyses. The statistical models for the analysis of metabolite profiles accounted for various experimental effects as detailed by de Abreu e Lima et al. (2017). After applying quality checks and computing BLUEs, 92 leaf metabolic analytes and 283 root metabolic analytes remained for further analyses. BLUEs for transcriptomic data were computed using the R-package *limma* (Ritchie, Phipson, Wu, Hu, Law, Shi and Smyth, 2015) in reference to established protocols (Smyth and Speed, 2003, Ritchie et al., 2007, Frisch et al., 2010). The design matrix for the linear model was based on the dye-labeling of a reference genotype. To account for possible differences between the microarrays, replicates of some genotypes across at least two factorials were included, and modeled through a fixed effect term. All gene expression values were subsequently computed, based on the log-ratio relative to this common genotype (Smyth, 2004). In total, 1,323 gene expression profiles were available. Repeatabilities (*w*^2^) were estimated for each endophenotype at the inbred line level using the same models as for the computation of BLUEs, but treating the genotype effect as random. This analysis was performed jointly for the Dent and Flint lines allowing for different means and heterogeneous genotypic variances of the heterotic groups, but assuming a common error variance. Variance components were estimated by Gibbs sampling using the *R* package *MCMCglmm* (Hadfield, 2010).

### Prediction models and model evaluation

Predictions of hybrid performance were compared on the basis of the core set of hybrids *H* and the corresponding sets of parent lines *D* and *F* on which data for all five predictors (P, pedigree; G, genomic; T, transcriptomic; L, leaf metabolic; R, root metabolic data) were available with the exception of data on a few lines missing at random for R due to fungal contamination. The matrices **W**_*D*_ and **W**_*F*_ are matrices of standardized feature measurements for the various predictors (G, T, L, R). The matrix **W** has dimension ’number of parent lines in the corresponding heterotic group’ (*n*_*D*_ = 142, *n*_*F*_ = 103) times ’number of features’ (*w*_*G*_ = 21, 565, *w*_*T*_ = 1, 323, *w*_*L*_ = 92, *w*_*R*_ = 283). The columns in **W**_*D*_ and **W**_*F*_ are centered and standardized to unit variance, respectively.

The kernels pertaining to each predictor and lines from each heterotic group — corresponding to genomic relationship matrices in the case of SNPs — can then be defined as

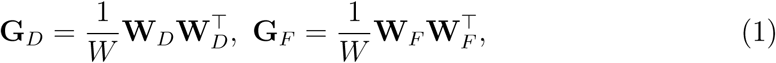

where *W* denotes the number of features (VanRaden, 2008). In the case of pedigree data (P), coancestry coefficients were used directly for **G**_*D*_ and **G**_*F*_, respectively.

The universal model for GCA and SCA effects was:

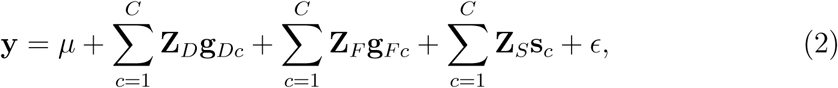

where **y** is the vector of observed hybrid performance (BLUEs), *μ* is the fixed model intercept, **Z**_*D*_ is the corresponding design matrix associating the random GCA effects of the lines in *D* (**g**_*Dc*_) with **y**, **Z**_*F*_ is the corresponding design matrix associating the random GCA effects of the lines in *F* (**g**_*Fc*_) with **y** and **Z**_*S*_ is a design matrix associating the SCA effects (**s**_*c*_), pertaining to hybrid combinations for the *c*-th predictor data type with the corresponding hybrid measurements in **y**. Thus, the model in Eq. 2 can accomodate just one (*C* = 1) or multiple (*C >* 1) predictors simultaneously. The random effects (**g**_*Dc*_ and **g**_*Fc*_) have expectation zero and covariance matrices equal to 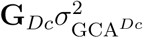 and 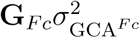 for the GCA effects of the Dent and Flint lines, respectively, **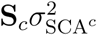** for the SCA effects and 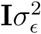 for the residual error. For each combination between crosses of lines *i* × *k* and *j × l*, the corresponding elements in **S**_*c*_ were obtained as the product of the respective elements *f*_*ij*_ in **G**_*Dc*_ and *f*_*kl*_ in **G**_*Fc*_, respectively (Schnell, 1965, Henderson, 1985, Bernardo, 1996, Massman et al., 2013, Technow et al., 2014, Jiang and Reif, 2015) (File S1). Note that, in the majority of cases, only GCA effects were considered. In the absence of epistasis, this model is equivalent to a feature model accounting for dually defined additive effects in each heterotic group and dominance effects between them. Extensions of the single-predictor models were made by adding GCA and SCA effects for any additional predictor assuming stochastic independence of effects. In order to obtain unbiased estimates of the predictive ability and to compare different models and predictor combinations, following Technow et al. (2014), we devised a cross-validation (CV) scheme, stratified by the parent lines and using 1,000 runs (CV1000, File S1). All prediction models were implemented using the *R* package *BGLR* (Pérez and de Los Campos, 2014).

### Comparison of predictive abilities

Predictive abilities were obtained by calculating Pearson correlations between predicted (*ŷ*) and observed phenotypes (*y*), separately for three test set partitions (T0, T1 and T2 hybrids). For each CV run, the training and validation sets were stored to ensure the validity of comparisons between any predictor and combinations thereof. For any two predictors, say *A* and *B*, we then have orthogonal vectors with predictive abilities *r*_*A*_ and *r*_*B*_ of length ’number of cross validation runs’.

### Evaluation of a pre-selection bias in transcriptomic data

A custom 2K-microarray (GPL22267) was assembled from a subset of the 47K maize oligonucleotide array (GPL6438), based on association of genes with hybrid performance or mid-parent heterosis for grain yield and grain dry matter content of maize. These two traits were evaluated in separate grain-yield trials with hybrids from factorial *H*_FAC^(1)^_ (Frisch et al. (2010), Thiemann et al. (2010), File S1). To ensure that no pre-selection bias was introduced in hybrid prediction using these transcriptomic data, we compared predictive abilities among the various predictors when excluding *H*_FAC^(1)^_ from the entire set *H*.

### Association mapping

For each of the seven agronomic traits, we performed a genome-wide association study (GWAS) with GCA effects of all 142 Dent and 103 Flint parent lines as response variables using the EMMAX-method (Kang et al., 2010) as implemented in *cpgen* (Heuer, 2015). To avoid using the marker data twice, GCA effects were calculated using only pedigree information. Furthermore, an eQTL analysis was carried out to examine statistically significant associations between genomic and transcriptomic data for the parent lines (*D* and *F*) of the core set *H* plus five additional lines. This was accomplished in the same way as in the GWAS for agronomic traits, but here the BLUPs of the transcriptomic data of each mRNA were used as the response variables. Associations in each GWAS were declared statistically significant at *α* = 0.05 after Bonferroni correction.

### Probability of success

Following Robson, Powers and Urquhart (1967), we calculated the probability of success (*P* [*r, β*]) that a hybrid, selected at random from the upper *β* percent fraction of the distribution of predicted values for predictor A, has a phenotypic value contained in the upper *β* percent of the distribution of observed values. Denoting the predictive ability of a given predictor by *r*, this conditional probabilty was calculated assuming a bivariate normal distribution

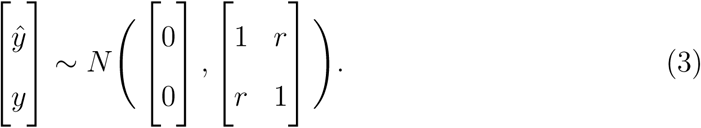

The required integrals were solved within the *R* statistical environment using the *mvtnorm* package (Genz et al., 2017).

### Principal component analysis

Principal component analyses (PCA) were carried out to examine whether different predictors can distinguish between Dent and Flint parent lines and to explore whether subpopulations exist within either hetrotic group. Prior to each PCA, all variables were scaled and centered. Clusters represent two component mixtures of bivariate t-distributions, which were estimated using Maximum Likelihood. Ellipses were drawn based on the 0.95 quantiles of the respective bivariate t-distributions. Unless stated otherwise, all statistical analyses were carried out inside the *R* environment for statistical computing (R Core Team, 2016).

### Data availability

The data and the code used to analyze the data are available upon request.

## Results

### Agronomic data

Mean values of the 1,536 hybrids for the seven evaluated agronomic traits, relevant for animal feed and biogas production, were of the same magnitude as reported by Riedelsheimer et al. (2012) and Grieder et al. (2011). For all traits, 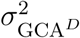 and 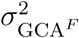, describing the main effects of the parents from each heterotic group, together explained more than 93% of the genotypic variance among hybrids (Table 1). Heritabilities were moderate to high for all agronomic traits, indicating a high precision of field experiments and data collection (Table 1).

**Table 1:**
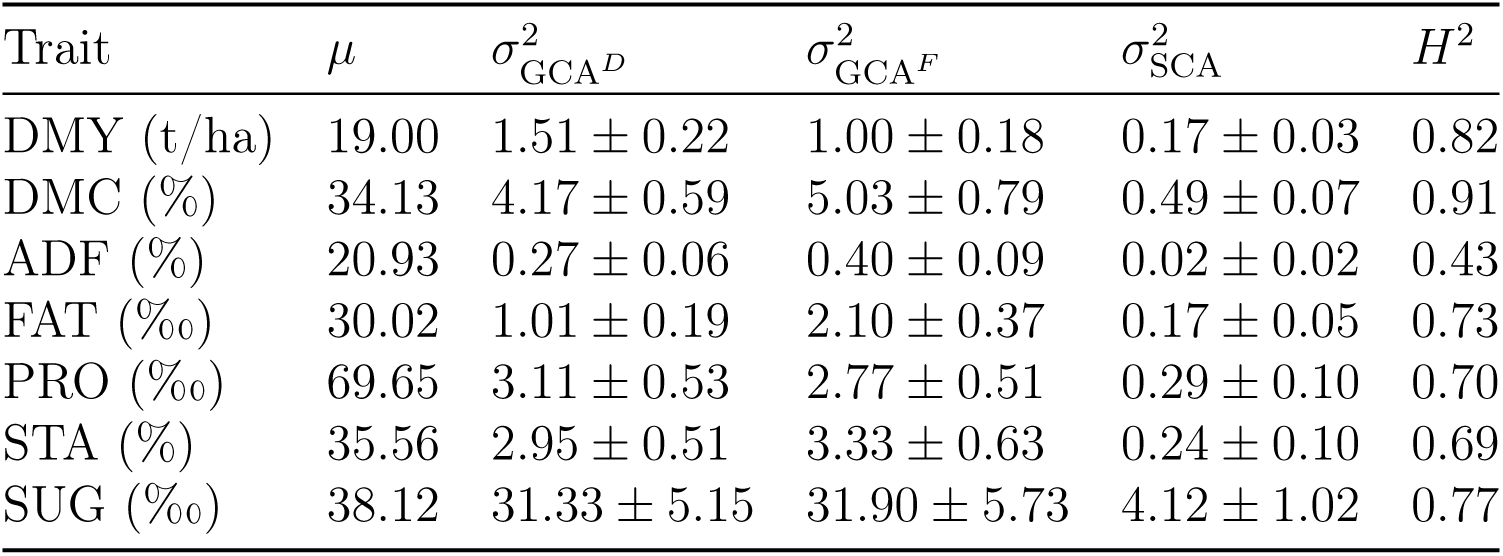
Summary of agronomic traits. Traits are characterized by overall mean (*μ*), variance components of GCA effects for Dent (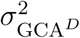) and Flint lines (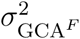) and SCA effects (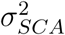) (followed by s.e.m.) as well as entry mean heritabilities (*H*^2^).

| Trait | $\mu$ | $\sigma^2_{\text{GCA}^D}$ | $\sigma^2_{\text{GCA}^F}$ | $\sigma^2_{\text{SCA}}$ | $H^2$ |
| --- | --- | --- | --- | --- | --- |
| DMY (t/ha) | 19.00 | $1.51 \pm 0.22$ | $1.00 \pm 0.18$ | $0.17 \pm 0.03$ | 0.82 |
| DMC (%) | 34.13 | $4.17 \pm 0.59$ | $5.03 \pm 0.79$ | $0.49 \pm 0.07$ | 0.91 |
| ADF (%) | 20.93 | $0.27 \pm 0.06$ | $0.40 \pm 0.09$ | $0.02 \pm 0.02$ | 0.43 |
| FAT (‰) | 30.02 | $1.01 \pm 0.19$ | $2.10 \pm 0.37$ | $0.17 \pm 0.05$ | 0.73 |
| PRO (‰) | 69.65 | $3.11 \pm 0.53$ | $2.77 \pm 0.51$ | $0.29 \pm 0.10$ | 0.70 |
| STA (%) | 35.56 | $2.95 \pm 0.51$ | $3.33 \pm 0.63$ | $0.24 \pm 0.10$ | 0.69 |
| SUG (‰) | 38.12 | $31.33 \pm 5.15$ | $31.90 \pm 5.73$ | $4.12 \pm 1.02$ | 0.77 |

### Predictor data

Repeatabilities (*w*^2^) for endophenotypes varied considerably in both groups of parents (Fig. S1a) with average values ranging from 0.31 to 0.41, except for transcriptomic data in Flint material where the average repeatability was only 0.18. Nevertheless, in the latter case, 291 out of 1,323 transcripts still exceeded a threshold of *w*^2^ = 0.4.

Dent and Flint lines were clearly separated in principal component analyses of genomic and transcriptomic data (Fig. 1a) without signs of subpopulations within either group. However, they overlapped for leaf metabolic and, to an even greater extent, for root metabolic data. Off-diagonal elements of the kernels **G**_*D*_ and **G**_*F*_, respectively, showed moderate correlations between genomic and transcriptomic data (*ρ*_*D*_ *≈* 0.56, *ρ*_*F*_ *≈* 0.44, Fig. S2). Correlations between the off-diagonal elements of the **G**-matrices were highest for the comparison between genomic and pedigree data (*ρ*_*D*_ *≈* 0.72, *ρ*_*F*_ *≈* 0.63). Intriguingly, the associations between the **G**-matrices for the root and leaf metabolic data were very low (*ρ*_*D*_ *≈* 0.12, *ρ*_*F*_ *≈* 0.06).

**Figure 1:**
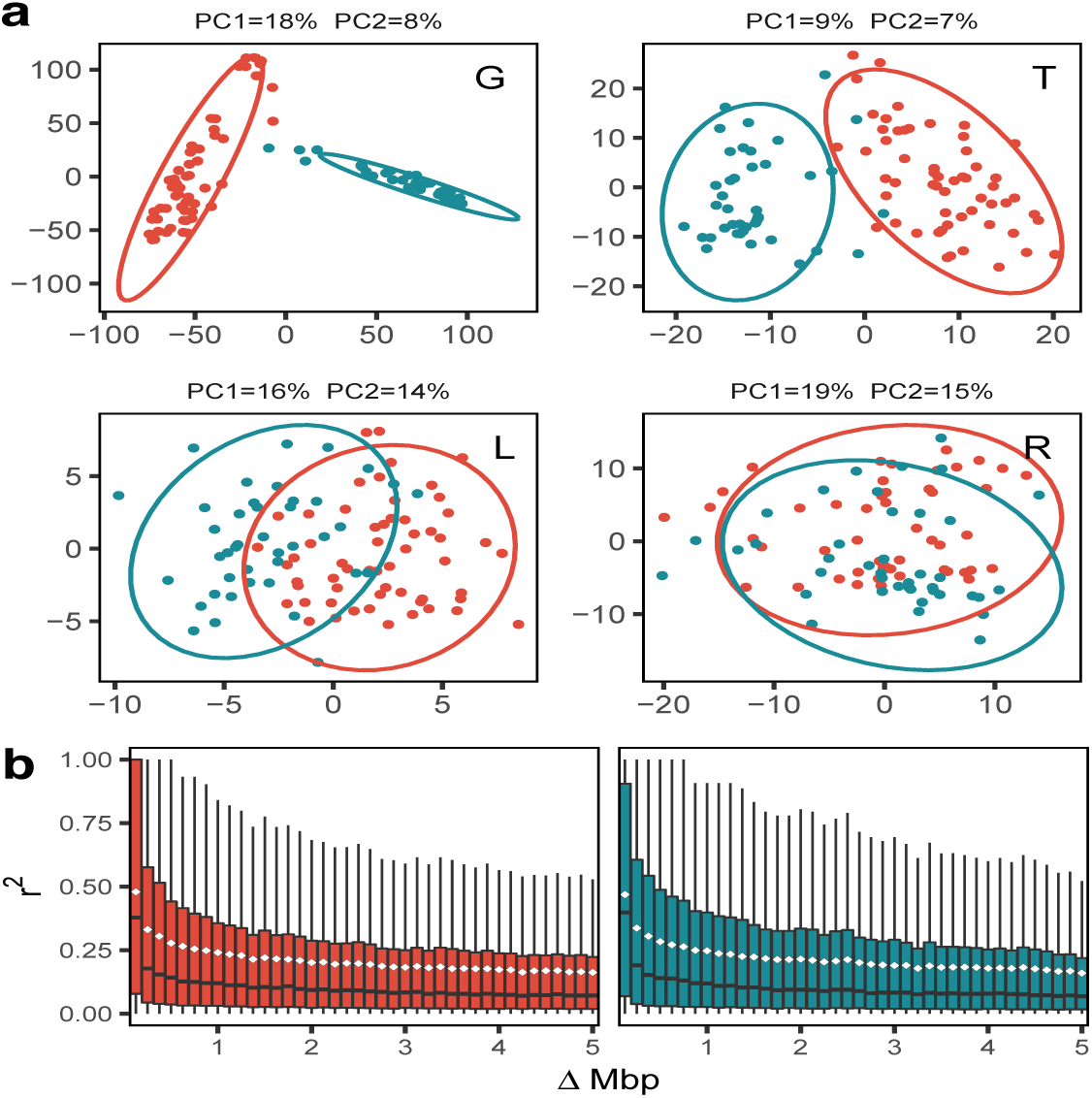
Properties of predictor data for Dent (red) and Flint (teal) parent lines. (**a**) Principal component analysis (PCA) of both groups for genomic (G), transcriptomic (T), leaf metabolic (L) and root metabolic (R) data. The variance explained by PC 1 (x axis) and PC 2 (y axis) are shown in the caption of each facet. (**b**) Linkage disequilibrium decay as a function of the distance between two loci using 40 bins of 0.125 Mbp width, each. The median *r*^2^ is depicted as a horizontal bar whereas the mean *r*^2^ is depicted as a white diamond.

We observed high median pairwise linkage disequilibrium (LD) between SNP markers (*r*^2^ *≈* 0.39 in Dent and *r*^2^ *≈* 0.37 in Flint material) at a distance of ΔMbp *≤* 0.125 (Fig. 1b). After an initial drop in *r*^2^ for Δ *>* 0.125, substantial long-range LD remained. Large differences in allele frequencies in the two heterotic groups were present for 57% of SNPs (Fig. 2a,b) — particularly in the telomeric regions of the genome. An eQTL analysis performed with the parent lines suggests that transcript abundance integrates variegated genetic information given the fact that i) on the same chromosome, significant associations not only occurred between adjacent but also between distant pairs of expressed genes and SNPs and ii) 50% of the significant associations (*α* = 0.05, Bonferroni-corrected) occurred between expressed genes and SNPs on different chromosomes (Fig. 2).

**Figure 2:**
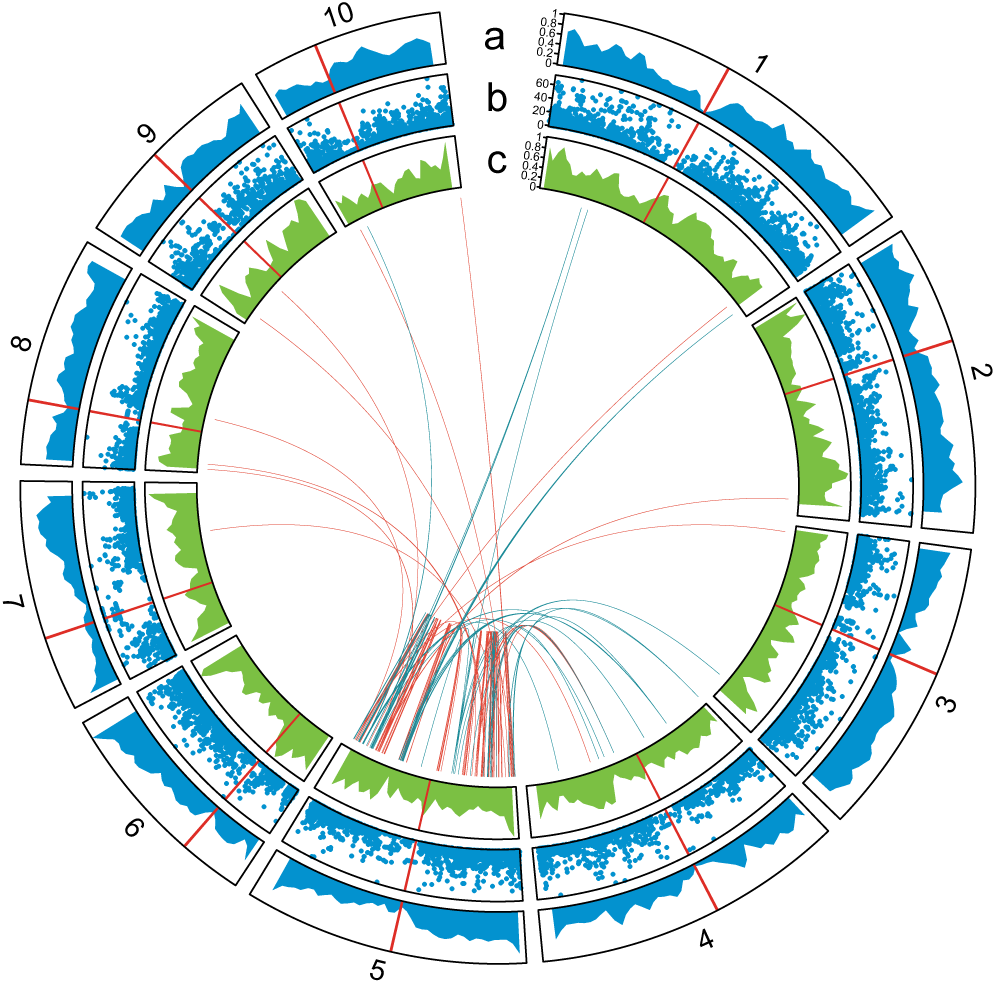
Distribution and relationship of genomic and transcriptomic data for the ten maize chromosomes. Centromeres for each chromosome are depicted as vertical red lines. (**a,c**) Density of SNPs and mRNAs, respectively, across the ten maize chromosomes. (**b**) Statistical significance (– log_10_ p-value) for differences in SNP allele frequencies between Dent and Flint lines. (**Center**) Links between any statistically significant (*α* = 0.05 after Bonferroni correction) association between SNPs and mR-NAs. Associations are displayed as links for SNPs on chromosome 5, for which the distribution of associations is representative for the entire genome, using red color for Dent parent lines and teal color for Flint parent lines.

### Predictive abilities

Assuming a polygenic architecture for all traits, as suggested by results from a GWAS (Fig. S3), we chose the best linear unbiased predictor (BLUP) method as a baseline for prediction of T0, T1 and T2 hybrids. Given that we corrected for fixed effects in advance, this method corresponds to a selection index. A cross-validation scheme with 1,000 runs (CV1000), stratified by the parent lines, was devised (File S1, Fig. S4). Our main emphasis was on predicting T0 hybrids given the fact that they constitute the majority of possible hybrids in practical breeding programs (Kadam et al., 2016).

For predictive abilities (*r*) of T0 hybrids, transcriptomic data alone were the best predictor for the most complex and highly heterotic trait, DMY, as well as for PRO (Fig. 3a). With transcriptomic data, the predictive ability *r* for DMY was 14.9% higher than for genomic data, resulting in an 85% increase in the probability of successfully selecting the best hybrid candidates *P* [*r, β*] for *β* = 0.01% (Fig. 3b). This selection intensity corresponds to picking the top 100 out of 10^6^ predicted hybrids for production and intensive testing in field trials.

**Figure 3:**
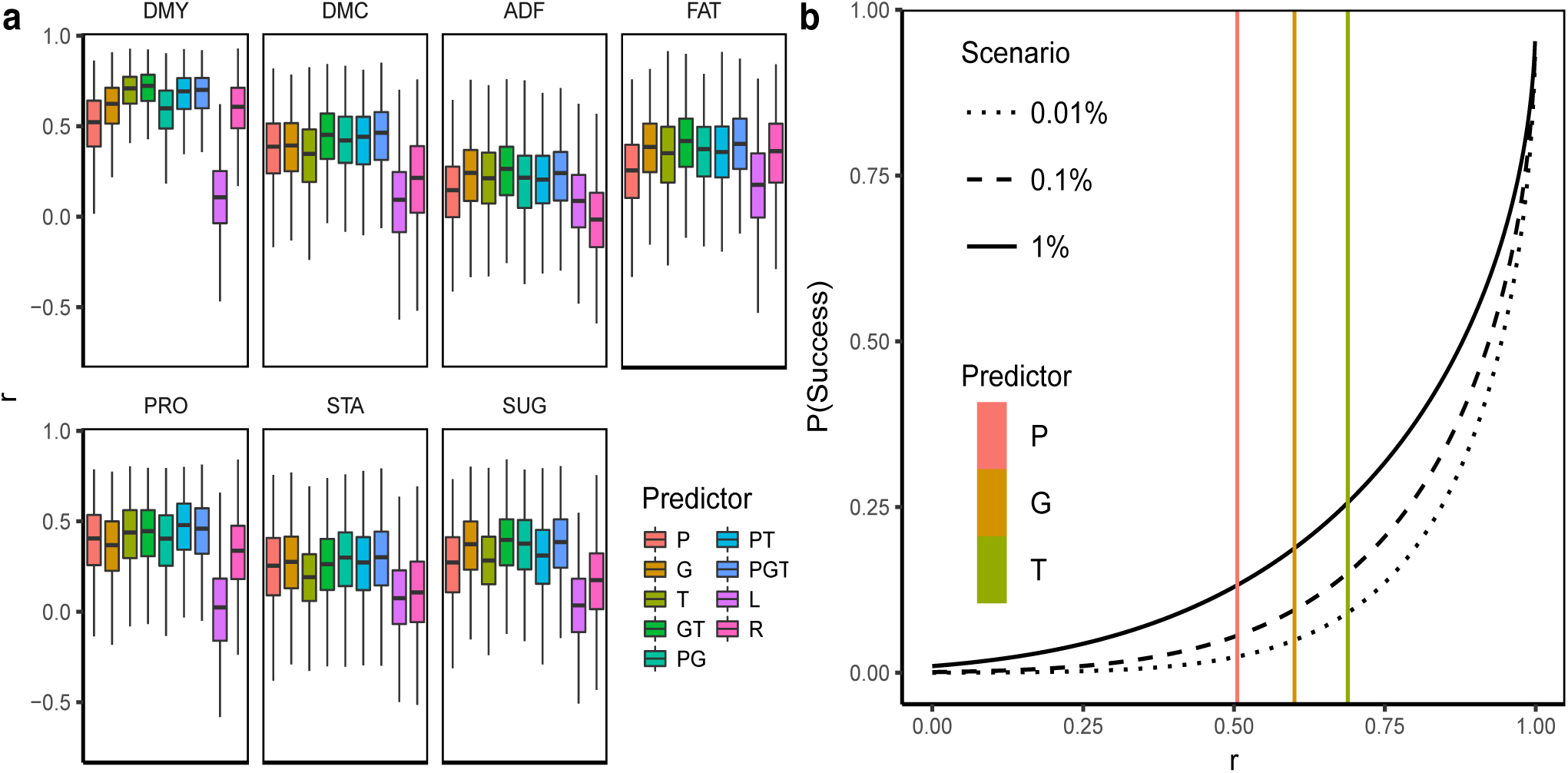
Predictive abilities (r) from BLUP models using a CV scheme with sampling of *|H*_*TRN*_*|* = 200 hybrids, *|D*_*TRN*_*|* = 40 Dent and |*F*_*TRN*_| = 33 Flint parent lines for various predictors and combinations thereof (P, pedigree; G, genome; T, transcriptome; L, leaf metabolome; R, root metabolome). (**a**) Comparison of *r* values from 1,000 CV runs for T0 hybrids and seven agronomic traits. (**b**) Success rate of selecting superior hybrids (*P* [*r, β*]). *P* [*r, β*] is a function of the predictive ability *r* = *r*(*y, ŷ*) and refers to the conditional probability of a hybrid, selected at random from the upper *β*% fraction of the distribution of predicted values (*ŷ*), having a phenotypic value contained in the upper *β*% of the distribution of phenotypic values *y*. Observed predictive abilities (*r*) for T0 hybrids and the trait DMY are displayed as vertical, colored lines for three predictors.

Compared to other individual predictors, *r* obtained with genomic data alone were higher for FAT and SUG. Root metabolites displayed moderate to high predictive abilities for DMY and FAT, but did not perform well otherwise. Leaf metabolites performed relatively poorly for all traits. Regardless of the trait, combinations of genomic and transcriptomic information displayed robust and consistently high predictive abilities. Except for PRO, incorporating additional endophenotypes as predictors into our models did not yield notable improvements but remained at the same level compared to combining genomic and transcriptomic data. Incorporating SCA effects into our models did not further improve predictive abilties (Fig. S5). Results for the combination of other predictors with metabolic data are not presented because no improvement of predictive abilities over the combination of genomic data with transcriptomic data and pedigree data could be achieved. Finally, we assessed the influence of the number of SNPs and mRNAs on predictive abilities. For genomic data, a subset of 5,000 SNPs already yielded the same predictive ability as when using the entire available set. For transcriptomic data, the predictive ability improved only marginally with subsets larger than 50% of the available transcripts (Fig. S6).

## Discussion

### A paradigm shift in hybrid breeding

Hybrid breeding programs are generally based on genetically divergent heterotic groups. Their use enables a better exploitation of heterosis when conducting crosses between them (Melchinger and Gumber, 1998) and is expected to reduce the ratio of specific to general combining ability variance (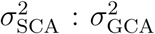) in the crosses, thereby allowing for the selection of hybrids largely on the basis of GCA of their parent lines (Reif et al., 2007). However, obtaining accurate estimates of GCA requires the evaluation of new lines in combinations with testers from the opposite heterotic group in multi-environment field trials. The promise of hybrid prediction is to accelerate breeding programs by skipping a large share of these tests in favor of selecting the most promising hybrids before they are even produced (Technow et al., 2014). This approach involves the prediction of an impressive number of putative hybrid candidates (*n*^2^) using predictor data collected on only 2*n* parent lines. Crucial for hybrid prediction are predictors, which not only reflect the relationship between parental inbred lines but also the interaction of the two parental genomes in their hybrid progeny.

#### Heterotic groups

Because of genetic drift and selection for hybrid performance, allele frequencies are expected to diverge in the two heterotic groups, thereby en-larging their genetic distance (Falconer and Mackay, 1996, Reif et al., 2007, Larièpe et al., 2017). Consistent with this hypothesis and two pilot studies with U.S. maize lines (Gerke et al., 2015, Hall et al., 2016), Dent and Flint lines in our study were clearly separated in principal component analyses of genomic and transcriptomic data. With large differences in allele frequencies *p*^*I*^ and *p*^*II*^ in the two heterotic groups, as observed for 57% of SNPs, dominance variance 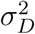 becomes very small because it is a function of the product *p*^*I*^(1 − *p*^*I*^)*p*^*II*^(1 − *p*^*II*^) (Stuber and Cockerham (1966), File S1).

Dominance variance (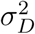) is the main component contributing to the variance of the specific combining ability effects (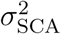), describing all types of interactions among the parental genomes in hybrid combinations. It was therefore not surprising that the variances of the general combining ability (GCA) effects (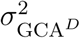 and 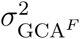), describing the main effects of the parents from each heterotic group, together explained more than 93% of the genotypic variance among hybrids for agronomic traits, which is consistent with earlier studies on silage maize of the Dent *×* Flint heterotic pattern (Geiger, Melchinger and Schmidt, 1986, Argillier, Méchin and Barrière, 2000). While the magnitude of SCA effects was trait specific, it was low for all observed traits, which is in agreement with previously reported values for yield and quality traits in silage maize (Grieder et al., 2012). The importance of GCA in our material was further corroborated by merely marginal differences in predictive abilities between models using only GCA effects and those that additionally incorporated SCA effects (Fig. S5). Nevertheless, in crops such as wheat, with yet no clearly defined heterotic groups (Zhao et al., 2015) and greater importance of SCA, inclusion of SCA effects in the model should improve predictive abilities.

#### Properties of well-established predictors

While pedigree data reflect the expected relationship between genotypes, they do not necessarily depict their realized relationship. Genomic data and downstream endophenotypes offer to improve upon this pedigree-based approximation by more closely mirroring the transmission of genes between genotypes and their interactions. Genomic data have the advantage of reliably capturing Mendelian sampling, thereby improving pedigree-based prediction for many traits. However, genomic data alone may not be the final answer for the prediction of complex traits for two major reasons: First, the number of samples in most studies is considerably smaller than the number of genetic markers or even nucleotides of a genome. This implies that just modeling additive effects already necessitates shrinkage of effects. More importantly, however, interactions between loci throughout the genome can be frequent (Brem et al., 2005, Brown et al., 2014), but attempts to incorporate this epistasis for the prediction of heterotic traits using genomic data have been disappointing when the prediction and training set did not share the same or closely related parents (Jiang and Reif, 2015). This was true even when using recently developed, efficient models (Jarquín et al., 2014, Martini et al., 2016) and suggests that genomic data capture only statistical epistasis, referring to genetic variation at the population level (Sackton and Hartl, 2016), which is generally of negligible magnitude (Hill, Goddard and Visscher, 2008, Mackay, 2014, Guo et al., 2016, Vazquez et al., 2016).

### Complementation of predictors

#### Flow of biological information

It is well-known that genetic effects on the phe-notype are mediated through multiple layers of endophenotypes (Civelek and Lusis, 2014, Ritchie, Holzinger, Li, Pendergrass and Kim, 2015) with information mainly flowing from the genome toward the phenotype via the transcriptome, the proteome and the metabolome with metabolite fluxes ultimately governing energy production and growth (Fiévet, Dillmann and de Vienne, 2010). For most traits in our material, metabolite- and pedigree-based predictive abilities were lower than those obtained with either transcriptomic or genomic information. However, consistently high predictive abilities across multiple traits could be realized when combining multiple predictors, as has been reported previously in humans (Vazquez et al., 2016) and maize inbred lines (Guo et al., 2016). This suggests complementary properties of the different predictors resulting in better proxies for the complex interplay in gene networks than genomic information alone. Such an advantage is particularly important for hybrid prediction when parents of prediction set hybrids are not closely related to parents of training set hybrids (File S1) as was shown by the relative excellence of transcriptomic data and the use of multiple predictors for the prediction of traits in T0 hybrids compared to T1 and T2 hybrids.

#### Tapping new sources of information

Whereas pedigree and genomic information are static, subsequent endophenotypes are characterized by pervasive interactions among and between each other (Dalchau et al., 2011, Zhu et al., 2012) and are, to varying degrees, influenced by biotic (Rudd et al., 2015, Tzin et al., 2015) and abiotic perturbations (Caldana et al., 2011, Witt et al., 2012). So while endophenotypes do not exclusively report on physiological epistasis but also on non-heritable effects, they seem to capture important information not represented by the genome given their intermediate position in the genotype-phenotype cascade. We get support for this hypothesis from (i) merely low to moderate correlations between off-diagonal elements of the kernels of different predictors in our study, (ii) mounting evidence for further improvements of predictive abilities when complementing genomic prediction with other endophenotypes despite sufficient marker densities (Fig. S6, Guo et al. (2016)) and (iii) the integration of SNP information from close and distant eQTL in the transcripts analyzed in our study. However, we concede that the number of parental genotypes in our mRNA assays was too small to warrant a reasonable statistical power for detecting epistasis in the expression of transcripts. In breeding programs, predictive abilities are largely driven by relationships — including Mendelian sampling — among genotypes compared to LD between SNP markers and causal QTL (Schopp et al., 2017). Increasing marker densities therefore have limited utility for improving genomic predictions as observed in our material, where SNP-based predictive abilities reached a plateau after using 5,000 equally spaced SNPs (Fig. S6). While two other studies also attempted to model interactions between different predictors, we refrained from this approach given that their reported predictive abilities based on interactions were not different from those in additive models despite using much larger sample sizes (Vazquez et al., 2016, Guo et al., 2016).

### Transcriptomic data

#### Utility of transcriptomic data for trait predictions

Of particular note was the excellent performance of transcriptomic data in predicting dry matter yield and protein. Evidence that parental gene expression patterns might be predictive of hybrid performance is given by (i) prevailing additive expression patterns in maize hybrids (Springer and Stupar, 2007*a*, Stupar et al., 2008), (ii) a positive correlation of the proportion of additive gene expression with the yield of hybrids (Guo et al., 2006), and (iii) co-localization of additively expressed genes with heterotic QTL (Thiemann et al., 2014). According to metabolic flux theory, gene expression in hybrids at the mid-parent level can generate hybrid vigor by counterbalancing opposing detrimental expression levels in their parent lines on a genome-wide scale (Kacser and Burns, 1981, Springer and Stupar, 2007*b*). The same concept is expected to apply to other quantitative endophenotypes (Lisec et al., 2011).

#### Pre-selection bias

As pointed out earlier, our transcripts were pre-selected based on associations with grain dry matter yield and grain dry matter content in hybrids, using a subset of the data included in our study (*H*_FAC^(1)^_). Hence, genotypes used for the pre-selection (*i.e. H*_FAC^(1)^_) could be regarded as a training set. By combining this “training set” and genotypes from the remaining five factorials, we might have introduced a bias by using predictors that have already seen the response variable in the *H*_FAC^(1)^_ genotypes. To rule out the existence of such a bias, we have compared the predictive abilities of different predictors for the complement of *H*_FAC^(1)^_. Two findings indicate that no bias in the comparison of predictive abilities was introduced: (i) Relative differences in predictive abilities between transcriptomic and pedigree or genomic data did not change when excluding genotypes from *H*_FAC^(1)^_ from the data (Fig. S7) and (ii) transcriptomic data performed rather poorly in predicting dry matter content although this trait was also among the criteria for the pre-selection procedure. Finally, an independent study using RNA-Seq data for the prediction of traits in maize inbred lines also reported exceptionally good performance of transcriptomic data in the prediction of multiple yield-related traits (Guo et al., 2016).

### Relative excellence of predictors for different traits

#### Tissue and sampling time

Despite the great prospects of using endophenotypes for trait predictions, some aspects require careful consideration when using this approach. A particular challenge in endophenotype-based prediction efforts is the choice of a suitable tissue and sampling time. Tissue-related effects regarding gene expression were found in studies on humans (Yang et al., 2015, Mele et al., 2015, Searle et al., 2016) and *A. thaliana* (Schmid et al., 2005) and in maize hybrids with respect to metabolome composition and metabolite abundance (Witt et al., 2012). Moreover, the age of an organism can selectively influence the expression of genes as observed in studies on humans (Mele et al., 2015, Yang et al., 2015) and *C. elegans* (Vinuela et al., 2010, Francesconi and Lehner, 2014). The low correlations between the off-diagonal elements of the kernels calculated from root and leaf metabolites might therefore be a reflection of highly dynamic processes differing between tissues and during different developmental stages. Whereas root metabolic data and transcriptomic data were obtained from seedlings germinated in standard controlled conditions, leaf metabolic data were derived from field-grown plants at a much later developmental stage, thereby increasing the possibility of environmentally-induced modifications. One might hypothesize, that the choice of sampling time and tissue could influence the chances of successful trait prediction if such age- or tissue-dependent transcripts and metabolites are associated with a phenotypic or clinical trait.

#### Feature selection

Another explanation for trait-dependent excellence of any predictor might lie in the sampling of features. In this study, only a small subset of metabolites was sampled and even very recent technologies (Xu et al., 2016, Dan et al., 2016) capture only a fraction of the estimated set of metabolites (Fernie, 2007). Moreover, the smaller differences in metabolite levels between both heterotic groups (Fig. 1) were most likely not conducive to capturing basic components un-derlying complex heterotic traits. It is also possible that transcriptomic data are associated with more biological processes than metabolite data and better capture the genetic effects relevant for the prediction of T0 hybrids.

#### Prospects for metabolites

Previously observed moderate metabolite-based pre-dictive abilities for T1 hybrids (Riedelsheimer et al., 2012) were confirmed in our study (Fig. S8), but for the majority of traits, root metabolites reached only medium and leaf metabolites even lower predictive abilities when predicting T0 hybrids. Despite the aforementioned shortcomings of metabolites, they have shown to be intriguing predictors due to their phsyiological proximity to the phenotype, which provides information that is impossible to infer from DNA or proteins (Fernie and Stitt, 2012), as well as encouraging results from other studies (Guo et al., 2016, Dan et al., 2016). A recently introduced technology, allowing for live-measurements of small molecules in the blood of living and awake animals (Arroyo-Currás et al., 2017), might overcome the problem of poorly time-resolved snap-shots of some metabolites with extremely fast turnover rates (Arrivault et al., 2009) if modified to properly work in plants.

#### Predictor requirements

Besides improving upon predictions based on pedigree relationships by capturing Mendelian sampling, the widespread use of genomic information in trait prediction has been driven by the ease of its application. In order to compete with genomic data, other ’omics’ data therefore require the use of standardized sampling conditions to obtain large repeatabilities and the possibility of season-independent sample extraction from seeds, seedlings or young roots to achieve high throughput.

### Conclusions

The use of whole-genome information has considerably advanced trait prediction over traditional pedigree-based BLUP by incorporating previously unobservable Mendelian sampling. Combining variegated sources of information promises to capture complex interactions between genes and endophenotypes, leading to stable predictions across traits. Especially if an extremely small fraction of the candidates is selected from the millions of possible new hybrids from each breeding cycle, the success of forecasts is a strongly convex function of predictive ability (Fig. 3b). Therefore, considering endophenotypes could have a substantial effect on the success and economics of hybrid breeding. Given the anticipated technological improvements in RNA-Seq and metabolite profiling, as well as the forthcoming adoption of the DH-technology for many crops (Kelliher et al., 2017), a paradigm shift from exclusively genomic prediction models to more inclusive approaches seems imminent.

## Author Contributions

W.S. and A.E.M developed the lines and hybrids, W.S. and A.E.M designed the field experiments, T.A.S analyzed the agronomic and pedigree data, L.W., M.S., A.S., A.E.M and M.W. designed the metabolic experiments, A.S. conducted the metabolic experiments, S.S. designed the transcriptomic experiments, F.S. and A.T. conducted the transcriptomic experiments, M.W. analyzed the metabolic and transcriptomic data, M.W., T.A.S, G.T., C.H. and A.E.M. devised the prediction models, M.W., C.H. and T.A.S. implemented the prediction models and developed software, H.F.U. and A.E. contributed to the statistical analysis. M.W., A.E.M., S.S, G.T., Z.N. and C.C.S. wrote the manuscript.

## Acknowledgements

The authors acknowledge support by the state of Baden-Württemberg through bwHPC. This project was funded by the German Federal Ministry of Education and Research (BMBF) within the projects OPTIMAL (FKZ: 0315958B,0315958F), SYNBREED (FKZ: 0315528D) and by the German Research Foundation (DFG, Grants No. ME 2260/5-1 and SCHO 764/6-1). Financial support for M.W. was provided by the Fiat Panis foundation, Ulm, Germany. We thank the staff of the Agricultural Experimental Research station, University of Hohenheim, for excellent technical assistance in conducting the field experiments. We are indebted to the group of R. Fries from Technische Universität München for the SNP genotyping of the parent inbred lines, to X. Mi for his assistance in preparing auxiliary figures based on the Mathematica software, to C. Zenke for advice on the computation of transcriptomic BLUEs and to P. Schopp for advice on prediction models.

## References

Argillier, O., V. Méchin and Y. Barrière. 2000. “Inbred line evaluation and breeding for digestibility-related traits in forage maize.” Crop Sci. 40(6):1596–1600.

Arrivault, Stephanie, Manuela Guenther, Alexander Ivakov, Regina Feil, Daniel Vosloh, Joost T. Van Dongen, Ronan Sulpice and Mark Stitt. 2009. “Use of reverse-phase liquid chromatography, linked to tandem mass spectrometry, to profile the Calvin cycle and other metabolic intermediates in Arabidopsis rosettes at different carbon dioxide concentrations.” Plant J. 59(5):824–839.

Arroyo-Currás, Netzahualcóyotl, Jacob Somerson, Philip A Vieira, Kyle L Ploense, Tod E Kippin and Kevin W Plaxco. 2017. “Real-time measurement of small molecules directly in awake, ambulatory animals.” Proc. Natl. Acad. Sci. U.S.A. 114(4):645–650.

Bernardo, Rex. 1996. “Best Linear Unbiased Prediction of Maize Single-Cross Performance.” Crop Sci 36:50–56.

Brem, Rachel B, John D Storey, Jacqueline Whittle and Leonid Kruglyak. 2005. “Genetic interactions between polymorphisms that affect gene expression in yeast.” Nature 436(7051):701–3.

Brown, Andrew Anand, Alfonso Buil, Ana Vinuela, Tuuli Lappalainen, Hou Feng Zheng, John B. Richards, Kerrin S. Small, Timothy D. Spector, Emmanouil T. Dermitzakis and Richard Durbin. 2014. “Genetic interactions affecting human gene expression identified by variance association mapping.” eLife 2014(3):1–16.

Browning, Brian L and Sharon R Browning. 2009. “A unified approach to genotype imputation and haplotype-phase inference for large data sets of trios and unrelated individuals.” Am. J. Hum. Genet. 84(2):210–223.

Butler, D, B R, Cullis, A R Gilmour and B J Gogel. 2009. “Mixed models for S language environments.”.

Caldana, Camila, Thomas Degenkolbe, Alvaro Cuadros-Inostroza, Sebastian Klie, Ronan Sulpice, Andrea Leisse, Dirk Steinhauser, Alisdair R Fernie, Lothar Willmitzer and Matthew a Hannah. 2011. “High-density kinetic analysis of the metabolomic and transcriptomic response of Arabidopsis to eight environmental conditions.” Plant J. 67(5):869–84.

Civelek, Mete and Aldons J Lusis. 2014. “Systems genetics approaches to understand complex traits.” Nat. Rev. Genet. 15(1):34–48.

Cox, T. S., J. P. Murphy and D. M. Rodgers. 1986. “Changes in genetic diversity in the red winter wheat regions of the United States.” Proc. Natl. Acad. Sci. U.S.A. 83(15):5583–5586.

Dalchau, Neil, Seong J Baek, Helen M Briggs, Fiona C Robertson, Antony N Dodd, Michael J Gardner, Matthew A Stancombe, Michael J Haydon, Guy-Bart Stan, Jorge M Gonçalves and Alex A R Webb. 2011. “The circadian oscillator gene GIGANTEA mediates a long-term response of the Arabidopsis thaliana circadian clock to sucrose.” Proc. Natl. Acad. Sci. U.S.A. 108(12):5104–9.

Dan, Zhiwu, Jun Hu, Wei Zhou, Guoxin Yao, Renshan Zhu, Yingguo Zhu and Wenchao Huang. 2016. “Metabolic prediction of important agronomic traits in hybrid rice (Oryza sativa L.).” Nature Sci. Rep. 6(October 2015):1–9.

de Abreu e Lima, Francisco, Matthias Westhues, Lothar Willmitzer, Albrecht E Melchinger and Zoran Nikoloski. 2017. “Metabolic robustness in young roots underpins a predictive model of maize hybrid performance in the field.” Plant J. pp. 1–11.

Duvick, D N. 2005. “Genetic progress in yield of United States maize (Zea mays L).” Maydica 50:193–202.

Falconer, D.S. and Trudy F C Mackay. 1996. Introduction to Quantitative Genetics 4th ed. Essex: Pearson.

Fernie, Alisdair R. 2007. “The future of metabolic phytochemistry: Larger numbers of metabolites, higher resolution, greater understanding.” Phytochemistry 68(22-24):2861–2880.

Fernie, Alisdair R and Mark Stitt. 2012. “On the discordance of metabolomics with proteomics and transcriptomics: coping with increasing complexity in logic, chemistry, and network interactions scientific correspondence.” Plant Physiol. 158(3):1139–45.

Fiévet, Julie B, Christine Dillmann and Dominique de Vienne. 2010. “Systemic properties of metabolic networks lead to an epistasis-based model for heterosis.” Theor. Appl. Genet. 120(2):463–73.

Francesconi, Mirko and Ben Lehner. 2014. “The effects of genetic variation on gene expression dynamics during development.” Nature 505(7482):208–11.

Frisch, Matthias, Alexander Thiemann, Junjie Fu, Tobias a. Schrag, Stefan Scholten and Albrecht E. Melchinger. 2010. “Transcriptome-based distance measures for grouping of germplasm and prediction of hybrid performance in maize.” Theor. Appl. Genet. 120(2):441–450.

Ganal, Martin W, Gregor Durstewitz, Andreas Polley, Aurélie Bérard, Edward S Buckler, Alain Charcosset, Joseph D Clarke, Eva-Maria Graner, Mark Hansen, Johann Joets, Marie-Christine Le Paslier, Michael D McMullen, Pierre Montalent, Mark Rose, Chris-Carolin Schön, Qi Sun, Hildrun Walter, Olivier C Martin and Matthieu Falque. 2011. “A large maize (Zea mays L.) SNP genotyping array: development and germplasm genotyping, and genetic mapping to compare with the B73 reference genome.” PloS ONE 6(12):e28334.

Geiger, H. H., Albrecht E. Melchinger and G.A. Schmidt. 1986. Analysis of factorial crosses between flint and dent maize inbred lines for forage performance and quality traits. In Breeding of Silage Maize, ed. O Dolstra and P Miedema. Wageningen: Pudoc pp. 147–154.

Genz, Alan, Frank Bretz, Tetsuhisa Miwa, Xuefei Mi, Friedrich Leisch, Fabian Scheipl and Torsten Hothorn. 2017. “mvtnorm: Multivariate Normal and t Distributions.”. URL: http://cran.r-project.org/package/mvtnorm

Gerke, Justin P., Jode W. Edwards, Katherine E. Guill, Jeffrey Ross-Ibarra and Michael D. McMullen. 2015. “The genomic impacts of drift and selection for hybrid performance in maize.” Genetics 201(3):1201–1211.

Gottesman, Irving I. and Todd D. Gould. 2003. “The endophenotype concept in psychiatry: Etymology and strategic intentions.” Am J Psychiatry 160(4):636–645.

Grieder, Christoph, Baldev S. Dhillon, Wolfgang Schipprack and Albrecht E. Melchinger. 2012. “Breeding maize as biogas substrate in Central Europe: II. Quantitative-genetic parameters for inbred lines and correlations with testcross performance.” Theor. Appl. Genet. 124(6):981–988.

Grieder, Christoph, Greta Mittweg, B.S. Dhillon, J.M. Montes, Elena Orsini and Albrecht E. Melchinger. 2011. “Determination of methane fermentation yield and its kinitics by near infrared spectroscopy and chemical composition in maize.” J Near Infrared Spectrosc 19(6):463–477.

Guo, Mei, Mary a Rupe, Xiaofeng Yang, Oswald Crasta, Christopher Zinselmeier, Oscar S Smith and Ben Bowen. 2006. “Genome-wide transcript analysis of maize hybrids: allelic additive gene expression and yield heterosis.” Theor. Appl. Genet. 113(5):831–845.

Guo, Zhigang, Michael M. Magwire, Christopher J. Basten, Zhanyou Xu and Daolong Wang. 2016. “Evaluation of the utility of gene expression and metabolic information for genomic prediction in maize.” Theor. Appl. Genet. 129(12):2413–2427.

Hadfield, J D. 2010. “MCMC methods for multi-response generalized linear mixed models: the MCMCglmm R package.” J Stat Softw 33(2):1–22.

Hall, Bradford D., Richard Fox, Qu Zhang, Andy Baumgarten, Barry Nelson, Joe Cummings, Ben Drake, Debora Phillips, Kevin Hayes, Mary Beatty, Gina Zastrow-Hayes, Brian Zeka, Jan Hazebroek and Stephen Smith. 2016. “Comparison of genotypic and expression data to determine distinctness among inbred lines of maize for granting of plant variety protection.” Crop Sci. 56(4):1443–1459.

Henderson, Charles R. 1984. Applications of Linear Models in Animal Breeding Models. Guelph: University of Guelph.

Henderson, C.R. 1985. “Best Linear Unbiased Prediction of Nonadditive Genetic Merits in Noninbred Populations.” J Anim Sci 60:111–117.

Heuer, Claas. 2015. “cpgen: Parallelized Genomic Prediction and GWAS.”. **URL:** https://cran.r-project.org/package=cpgen

Hill, William G., Michael E. Goddard and Peter M. Visscher. 2008. “Data and theory point to mainly additive genetic variance for complex traits.” PLoS Genet. 4(2):1–10.

Jarquín, Diego, José Crossa, Xavier Lacaze, Philippe Du Cheyron, Joëlle Daucourt, Josiane Lorgeou, François Piraux, Laurent Guerreiro, Paulino Pérez, Mario Calus, Juan Burgueño and Gustavo de los Campos. 2014. “A reaction norm model for genomic selection using high-dimensional genomic and environmental data.” Theor. Appl. Genet. 127(3):595–607.

Jiang, Yong and Jochen C. Reif. 2015. “Modelling epistasis in genomic selection.” Genetics 201(2):759–768.

Kacser, H and James A Burns. 1981. “The molecular basis of dominance.” Genetics 97:639–666.

Kadam, Dnyaneshwar, Sarah Potts, Martin O Bohn, Alexander E. Lipka and Aaron Lorenz. 2016. “Genomic Prediction of Hybrid Combinations in the Early Stages of a Maize Hybrid Breeding Pipeline.” G3 6:3443–3453.

Kang, Hyun Min, Jae Hoon Sul, Susan K Service, Noah a Zaitlen, Sit-Yee Kong, Nelson B Freimer, Chiara Sabatti and Eleazar Eskin. 2010. “Variance component model to account for sample structure in genome-wide association studies.” Nat. Genet. 42(4):348–354.

Kelliher, Timothy, Dakota Starr, Lee Richbourg, Satya Chintamanani, Brent Delzer, Michael L. Nuccio, Julie Green, Zhongying Chen, Jamie McCuiston, Wenling Wang, Tara Liebler, Paul Bullock and Barry Martin. 2017. “MATRILIN-EAL, a sperm-specific phospholipase, triggers maize haploid induction.” Nature 542(7639):105–109.

Kerr, M K and Gary A Churchill. 2001. “Experimental design for gene expression microarrays.” Biostatistics 2(2):183–201.

Larièpe, A., L. Moreau, J. Laborde, C. Bauland, S. Mezmouk, L. Décousset, T. Mary-Huard, J. B. Fiévet, A. Gallais, P. Dubreuil and A. Charcosset. 2017. “General and specific combining abilities in a maize (Zea mays L.) test-cross hybrid panel: relative importance of population structure and genetic divergence between parents.” Theor. Appl. Genet. 130(2):403–417.

Lisec, Jan, Lilla Römisch-Margl, Zoran Nikoloski, Hans-Peter Piepho, Patrick Gi- avalisco, Joachim Selbig, Alfons Gierl and Lothar Willmitzer. 2011. “Corn hybrids display lower metabolite variability and complex metabolite inheritance patterns.” Plant J. 68(2):326–336.

Longin, C. Friedrich H, Xuefei Mi and Tobias Würschum. 2015. “Genomic selection in wheat: optimum allocation of test resources and comparison of breeding strategies for line and hybrid breeding.” Theor. Appl. Genet. 128(7):1297–1306.

Mackay, Trudy F C. 2014. “Epistasis and quantitative traits: using model organisms to study gene-gene interactions.” Nat. Rev. Genet. 15(1):22–33.

Mackay, Trudy F C, Eric A Stone and Julien F Ayroles. 2009. “The genetics of quantitative traits: challenges and prospects.” Nat. Rev. Genet. 10(8):565–77.

Martini, Johannes W R, Valentin Wimmer, Malena Erbe and Henner Simianer. 2016. “Epistasis and covariance: how gene interaction translates into genomic relationship.” Theor. Appl. Genet. 129(5):963–976.

Massman, Jon M, Andres Gordillo, Robenzon E Lorenzana and Rex Bernardo. 2013. “Genomewide predictions from maize single-cross data.” Theor. Appl. Genet. 126(1):13–22.

Melchinger, Albrecht E and Ramesh K Gumber. 1998. Overview of heterosis and heterotic groups in agronomic crops. In Concepts and breeding of heterosis in crop plants, ed. K.R. Lamkey and J.E. Staub. Madison: CSSA p. 16.

Mele, M., P. G. Ferreira, F. Reverter, D. S. DeLuca, J. Monlong, M. Sammeth, T. R. Young, J. M. Goldmann, D. D. Pervouchine, T. J. Sullivan, R. Johnson, A. V. Segre, S. Djebali, A. Niarchou, T. G. Consortium, F. A. Wright, T. Lappalainen, M. Calvo, G. Getz, E. T. Dermitzakis, K. G. Ardlie and R. Guigo. 2015. “The human transcriptome across tissues and individuals.” Science 348(6235):660–665.

Mrode, Raphael A. 2014. Linear Models for the Prediction of Animal Breeding Values. 3rd ed. Oxfordshire: CABI.

Patti, Gary J, Oscar Yanes and Gary Siuzdak. 2012. “Metabolomics: the apogee of the omics trilogy.” Nat. Rev. Mol. Cell Biol. 13(4):263–9.

Pérez, Paulino and Gustavo de Los Campos. 2014. “Genome-wide regression & prediction with the BGLR statistical package.” Genetics 198(october):483–495.

R Core Team. 2016. “R: A Language and Environment for Statistical Computing.”. URL: https://www.r-project.org/

Reif, J C, F.M Gumpert, S Fischer and A. E Melchinger. 2007. “Impact of inter-population divergence on additive and dominance variance in hybrid populations.” Genetics 176(3):1931–1934.

Riedelsheimer, Christian, Angelika Czedik-Eysenberg, Christoph Grieder, Jan Lisec, Frank Technow, Ronan Sulpice, Thomas Altmann, Mark Stitt, Lothar Willmitzer and Albrecht E Melchinger. 2012. “Genomic and metabolic prediction of complex heterotic traits in hybrid maize.” Nat. Genet. 44(2):217–20.

Ritchie, M. E., B. Phipson, D. Wu, Y. Hu, C. W. Law, W. Shi and G. K. Smyth. 2015. “Limma powers differential expression analyses for RNA-sequencing and microarray studies.” Nucleic Acids Res. 43(7):e47.

Ritchie, Marylyn D., Emily R. Holzinger, Ruowang Li, Sarah A. Pendergrass and Dokyoon Kim. 2015. “Methods of integrating data to uncover genotype-phenotype interactions.” Nat. Rev. Genet. 16:85–97.

Ritchie, Matthew E, Jeremy Silver, Alicia Oshlack, Melissa Holmes, Dileepa Diyagama, Andrew Holloway and Gordon K Smyth. 2007. “A comparison of background correction methods for two-colour microarrays.” Bioinformatics 23(20):2700–2707.

Robson, D. S., LeRoy Powers and N. S. Urquhart. 1967. “The proportion of genetic deviates in the tails of a normal population.” Der Züchter Genetics and Breeding Research 37(4):205–216.

Rudd, Jason J, Kostya Kanyuka, Keywan Hassani-Pak, Mark Derbyshire, Ambrose Andongabo, Jean Devonshire, Artem Lysenko, Mansoor Saqi, Nalini M Desai, Stephen J Powers, Juliet Hooper, Linda Ambroso, Arvind Bharti, Andrew Farmer, Kim E Hammond-Kosack, Robert A Dietrich and Mikael Courbot. 2015. “Transcriptome and metabolite profiling of the infection cycle of Zymoseptoria tritici on wheat reveals a biphasic interaction with plant immunity involving differential pathogen chromosomal contributions and a variation on the hemibiotrophic lifestyle def.” Plant Physiol. 167(3):1158–85.

Sackton, Timothy B and Daniel L Hartl. 2016. “Perspective genotypic context and epistasis in individuals and populations.” Cell 166:279–287.

Schmid, Markus, Timothy S Davison, Stefan R Henz, Utz J Pape, Monika Demar, Martin Vingron, Bernhard Schölkopf, Detlef Weigel and Jan U Lohmann. 2005. “A gene expression map of Arabidopsis thaliana development.” Nat. Genet. 37(5):501–6.

Schnell, F.W. 1965. “Die Covarianz zwischen Verwandten in einer gen-orthogonalen Population. I. Allgemeine Theorie.” Biom Z 7(1):2–49.

Schopp, Pascal, Dominik Müller, Frank Technow and Albrecht E Melchinger. 2017. “Accuracy of genomic prediction in synthetic populations depending on the number of parents, relatedness and ancestral linkage disequilibrium.” Genetics 205:441–454.

Searle, Brian C, Rachel M Gittelman, Ohad Manor and Joshua M Akey. 2016. “Detecting Sources of Transcriptional Heterogeneity in Large-Scale RNA-Seq Data Sets.” Genetics 204(December):1391–1396.

Smyth, G.K. 2004. “Linear models and empirical bayes methods for assessing differential expression in microarrays experiments.” Stat Appl Genet Mol Biol 3(1):1–26.

Smyth, Gordon K and Terry Speed. 2003. “Normalization of cDNA microarray data.” Methods 31(4):265–273.

Springer, N. M. and R. M. Stupar. 2007a. “Allele-specific expression patterns reveal biases and embryo-specific parent-of-origin effects in hybrid maize.” Plant Cell 19(8):2391–2402.

Springer, Nathan M. and Robert M. Stupar. 2007b. “Allelic variation and heterosis in maize: How do two halves make more than a whole?” Genome Res. 17(3):264–275.

Stuber, C W and C Clark Cockerham. 1966. “Gene effects and variances in hybrid populations.” Genetics 54(6):1279–1286.

Stupar, Robert M., Jack M Gardiner, Aaron G Oldre, William J Haun, Vicki L Chandler and Nathan M Springer. 2008. “Gene expression analyses in maize inbreds and hybrids with varying levels of heterosis.” BMC Plant Biol. 8(33):1–19.

Swanson-Wagner, Ruth A, Yi Jia, Rhonda DeCook, Lisa A Borsuk, Dan Nettleton and Patrick S Schnable. 2006. “All possible modes of gene action are observed in a global comparison of gene expression in a maize F1 hybrid and its inbred parents.” Proc. Natl. Acad. Sci. U.S.A. 103(18):6805–6810.

Technow, Frank, Tobias A Schrag, Wolfgang Schipprack, Eva Bauer, Henner Simianer and Albrecht E Melchinger. 2014. “Genome properties and prospects of genomic prediction of hybrid performance in a breeding program of maize.” Genetics 197:1343–1355.

Thiemann, Alexander, Junjie Fu, Felix Seifert, Robert T Grant-Downton, Tobias a Schrag, Heike Pospisil, Matthias Frisch, Albrecht E Melchinger and Stefan Scholten. 2014. “Genome-wide meta-analysis of maize heterosis reveals the potential role of additive gene expression at pericentromeric loci.” BMC Plant Biol. 14(88):1–14.

Thiemann, Alexander, Junjie Fu, Tobias a. Schrag, Albrecht E. Melchinger, Matthias Frisch and Stefan Scholten. 2010. “Correlation between parental transcriptome and field data for the characterization of heterosis in Zea mays L.” Theor. Appl. Genet. 120(2):401–413.

Tzin, Vered, Noe Fernandez-Pozo, Annett Richter, Eric A Schmelz, Matthias Schoettner, Martin Schäfer, Kevin R Ahern, Lisa N Meihls, Harleen Kaur, Alisa Huffaker, Naoki Mori, Joerg Degenhardt, Lukas A Mueller and Georg Jander. 2015. “Dynamic Maize Responses to Aphid Feeding Are Revealed by a Time Series of Transcriptomic and metabolomic assays.” Plant Physiol. 169(november):1727–1743.

van den Berg, Robert A, Huub C J Hoefsloot, Johan A Westerhuis, Age K Smilde and Mariët J van der Werf. 2006. “Centering, scaling, and transformations: improving the biological information content of metabolomics data.” BMC Genomics 7:1–15.

VanRaden, P M. 2008. “Efficient methods to compute genomic predictions.” J. Dairy Sci. 91(11):4414–4423.

Vazquez, Ana I., Yogasudha C. Veturi, Michael Behring, Sadeep Shrestha, Matias Kirst, Marcio F.R. Resende Jr. and Gustavo de los Campos. 2016. “Increased proportion of variance explained and prediction accuracy of survival of breast cancer patients with use of whole-genome multi-omic profiles.” Genetics 203(3):1425–1438.

Vinuela, Ana, L Basten Snoek, Joost A G Riksen and Jan E Kammenga. 2010. “Genome-wide gene expression regulation as a function of genotype and age in C. elegans.” Genome Res. 20:929–937.

Wedzony, M., B.P. Forster, I. Zur, E. Golemiec, M. Scechynska-Hebda, E. Dubas and G. Gotebiowska. 2009. Progress in Doubled Haploid Technology in Higher Plants. In Advances in Haploid Production in Higher Plants, ed. Alisher Touarev, Brian P. Forster and Jain S. Mohan. Springer chapter 1, p. 33.

Witt, Sandra, Luis Galicia, Jan Lisec, Jill Cairns, Axel Tiessen, Jose Luis Araus, Natalia Palacios-Rojas and Alisdair R Fernie. 2012. “Metabolic and phenotypic responses of greenhouse-grown maize hybrids to experimentally controlled drought stress.” Mol. Plant 5(2):401–17.

Xu, Shizhong, Yang Xu, Liang Gong and Qifa Zhang. 2016. “Metabolomic prediction of yield in hybrid rice.” Plant J. 88(2):219–227.

Yang, Jialiang, Tao Huang, Francesca Petralia, Quan Long, Bin Zhang, Carmen Argmann, Yong Zhao, Charles V Mobbs, Eric E Schadt, Jun Zhu and Zhidong Tu. 2015. “Synchronized age-related gene expression changes across multiple tissues in human and the link to complex diseases.” Nat. Sci. Rep. 5:15145.

Zenke-Philippi, Carola, Alexander Thiemann, Felix Seifert, Tobias Schrag, Albrecht E. Melchinger, Stefan Scholten and Matthias Frisch. 2016. “Prediction of hybrid performance in maize with a ridge regression model employed to DNA markers and mRNA transcription profiles.” BMC Genomics 17(1):262.

Zhao, Yusheng, Zuo Li, Guozheng Liu, Yong Jiang, Hans Peter Maurer, Tobias Würschum, Hans-Peter Mock, Andrea Matros, Erhard Ebmeyer, Ralf Schachschneider, Ebrahim Kazman, Johannes Schacht, Manje Gowda, C Friedrich H Longin and Jochen C Reif. 2015. “Genome-based establishment of a high-yielding heterotic pattern for hybrid wheat breeding.” Proc. Natl. Acad. Sci. U.S.A. 112(51):15624–15629.

Zhu, Jun, Pavel Sova, Qiuwei Xu, Kenneth M. Dombek, Ethan Y. Xu, Heather Vu, Zhidong Tu, Rachel B. Brem, Roger E. Bumgarner and Eric E. Schadt. 2012. “Stitching together multiple data dimensions reveals interacting metabolomic and transcriptomic networks that modulate cell regulation.” PLoS Biol. 10(4).

